# Structure of the Type I-F3 CAST holo integration complex reveals licensing mechanisms during RNA-guided DNA integration

**DOI:** 10.64898/2026.07.01.735701

**Authors:** Shirin Fatma, Shubham Dubey, Vinh H. Truong, Seong Guk Park, Hailey M. Wallace, Alfredo Jose Florez Ariza, Isabella King, Elizabeth H. Kellogg

**Affiliations:** Structural Biology Department, St Jude Children’s Research Hospital, Memphis, TN; Center for Protein Design, Center of Excellence for Data Driven Discovery, St Jude Children’s Research Hospital, Memphis, TN; San Diego State University, San Diego, CA

## Abstract

Precise genomic integration of large DNA cargo remains a central challenge in genome engineering. In this context, CRISPR-associated transposases (CASTs) offer a promising solution by coupling RNA-guided DNA recognition to transposon-mediated integration. Type I-F3 CASTs exhibit exceptional targeting fidelity, making them a leading focus for biotechnology and gene therapy applications. Yet, despite extensive characterization, the structural basis for RNA-guided DNA integration of type I-F3 CASTs has remained elusive. Here, using cryo-EM and functional assays, we report the structural licensing mechanism of the *Vibrio cholerae* CAST (VchCAST) holo integration complex (HIC). Our structures reveal that sequential conformational licensing events govern selective activation of the integration machinery at the on-target site. Different components, including Cascade-TniQ, TnsC, and TnsA/B, undergo conformational changes upon assembly into the HIC, and the catalytically competent state is stabilized only within the fully assembled complex. Finally, comparison with existing CAST integration complex structures reveals a set of conserved architectural features, suggesting a common molecular basis for RNA-guided transposition across CAST diversity.

## Introduction

A longstanding challenge in genome engineering is how to achieve precise, programmable integration of large (≥1 kb) DNA cargo. Although CRISPR–Cas nucleases are capable of efficient DNA cleavage (*1*), their reliance on host DNA repair pathways limits both efficiency and fidelity of large insertions (*2*). CRISPR-associated transposases (CASTs) offer an alternative strategy by coupling RNA-guided DNA recognition to transposon-mediated integration, thus enabling targeted DNA insertion without the need for double-strand breaks or homologous recombination (*3*, *4*). CASTs evolved from Tn7-like transposons through the acquisition of a CRISPR effector, with distinct effectors defining each of the major subfamilies: type V-K, I-F3, I-B, and I-D (*5*). Despite this diversity, all CASTs encode a conserved set of transposition proteins: TniQ (bridging protein), TnsC (AAA+ regulator), and TnsA/B (transposase complex) (**Figure 1A**). The CRISPR effector and TniQ form the targeting module, while TnsA/B and TnsC form the transposition machinery (**Figure 1A**). A central mechanistic question in CAST biology is how this conserved transposition machinery has evolved to function with architecturally diverse CRISPR effectors.

**Figure 1.**
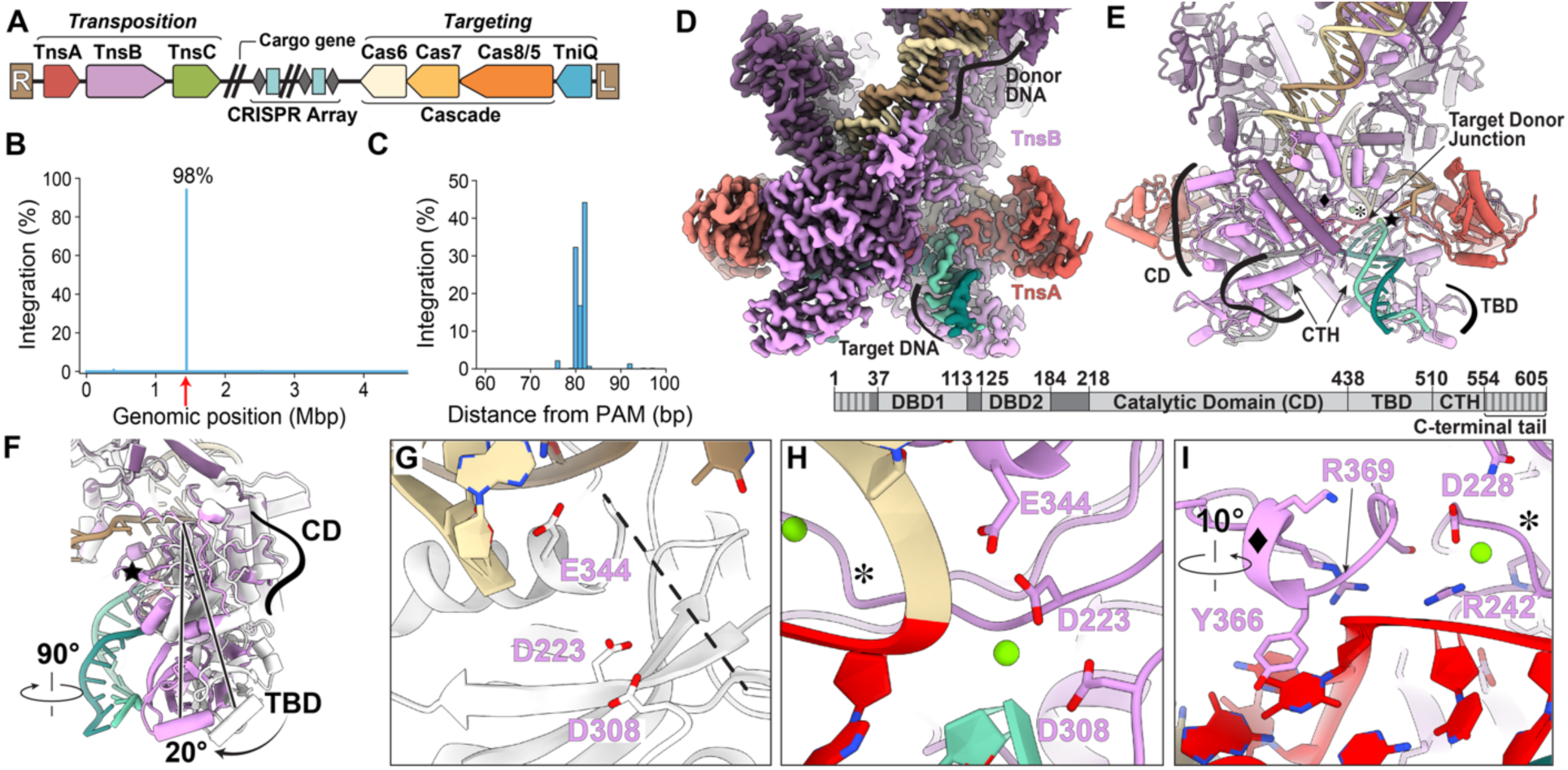
VchCAST exhibits highly specific RNA-guided DNA integration and reveals TnsA-dependent conformational changes promoting target DNA engagement. **(A)** Gene organization of the VchCAST transposon. R and L correspond to the right and left transposon ends, respectively. Genes are labeled and colored according to the convention adopted throughout this manuscript. Light blue boxes/grey diamonds indicate the CRISPR array. Hash marks indicate discontinuity. **(B)** *In vivo* genome-wide insertion profile of VchCAST. The x-axis indicates genomic position in megabase pairs (Mbp). The red arrow indicates the programmed target site. **(C)** Close-up view of the target site region shown in panel B, highlighting the insertion window downstream of the PAM motif. bp = base pair. **(D)** Cryo-EM density map (2.9 Å) of the strand-transfer complex (STC, Class 1), displayed at a threshold of 0.129. Each protein is colored as in panel A, with different shades to distinguish individual subunits. Donor DNA in shades of beige, target DNA in aqua, and the target site duplication in red. **(E)** Atomic model of the STC oriented as in panel D. Black symbols (asterisk, star, and diamond) indicate regions shown in panels F-I. TnsB domains are labeled in the model: catalytic domain (CD), target binding domain (TBD), and C-terminal helix (CTH). Bottom, domain organization of TnsB, with domain boundaries indicated by residue numbers; DBD1 and DBD2, DNA-binding domains 1 and 2, respectively. Striped regions indicate disordered N- and C-terminal tails. **(F)** Overlay of the catalytic TnsB in STC assembly Class 2 (white) and Class 1 (purple), from 90° rotated view with respect to panel E; star symbol placed in the same location as in panel E. The overlay shows conformational changes involved in transition from Class 2 to Class 1 models, with rotations of the catalytic domain (CD) and target binding domain (TBD) indicated with arrows and angles **(G)** Close-up view of the Class 2 model (white) showing the TnsB active site in the same orientation as panel E. Catalytic residues D223, D308, and E344 are shown as sticks. Dashed lines indicate unresolved residues: positions 331–341. **(H)** Same view as in panel I, showing the corresponding region in the Class 1 model. Catalytic residues are labeled, and the magnesium ion is shown as a green sphere. Asterisk corresponds to the same position indicated in panel E. **(I)** Close-up view of the region marked by the diamond and asterisk in panel E. Rotation is relative to the orientation shown in panel E. R242, D228, Y366, and R369 are shown in sticks and labeled.

While all CASTs perform RNA-guided transposition, they display a remarkable diversity of targeting behaviors. For example, in type I-B2 CASTs, the transposition machinery can associate with either the sequence-specific DNA-binding protein TnsD or an RNA–guided targeting complex (*6*). Type V-K CASTs exhibit a robust off-target pathway driven by the propensity of TniQ and TnsC to load at non-cognate sites (*7*). Type I-F3 CASTs have a Cascade effector and exhibit near-exclusive on-target integration (*3*), which makes them of particular interest for clinical (*8*) and microbial genome-editing applications (*9*).

Despite extensive biochemical, genetic, and structural studies of individual components, the mechanisms by which type I-F3 CASTs achieve precise on-target integration remain unknown. ChIP–seq experiments have revealed a hierarchical assembly pathway in which Cascade–TniQ occupancy precedes TnsC recruitment, and stable TnsC accumulation depends on the transposase TnsA/B (*10*). Although Cascade–TniQ binds DNA promiscuously and forms partial R-loops at many candidate sites (*10*), it appears insufficient to complete R-loop formation on its own (*11*), suggesting that additional steps are required to license TnsC recruitment. Structures of TnsC (*10*) have established the molecular details of its homo-oligomerization on DNA. Yet, how all these components converge to form an integration-competent complex at the RNA-guided target site remains unknown. Thus, a structure of the fully assembled holo integration complex is critical to establish the mechanistic basis for RNA-guided DNA integration and to guide rational engineering efforts to improve type I-F3 CAST integration efficiency for translational applications (*8*, *12*).

Here, using a combination of single-particle cryo-EM and directed *in vivo* mutagenesis, we uncover the mechanisms that regulate holo integration complex assembly in the type I-F3 *Vibrio cholerae* (VchCAST) system. The architectures of the strand-transfer complex (STC) and the full holo integration complex (HIC) reveal how RNA-guided target recognition, TnsC oligomerization, and TnsA/B recruitment converge to organize a stable integration complex assembly. Despite large structural differences across type V-K, I-F3, and I-B2 CASTs, conserved local features suggest that mechanistic principles of RNA-guided DNA integration may be universally shared. Our results reveal a context-dependent structural mechanism, where each of the aforementioned components undergoes significant conformational rearrangements, relative to their isolated structures (*10*, *11*, *13*), as a means to license RNA-guided integration complex assembly. The holo integration complex structure reported in this work should thus serve as a suitable blueprint for the rational engineering of highly active CAST variants.

## Results

### VchCAST exhibits highly specific RNA-guided DNA integration

The *Vibrio cholerae* CRISPR-associated transposase (VchCAST) comprises a conserved transposition machinery, consisting of TnsA, TnsB, and TnsC, coupled to a type I-F3 Cascade-TniQ targeting complex that directs RNA-guided DNA integration (**Figure 1A**). Previous studies established that *Vch*CAST integrates DNA at a defined distance downstream of the protospacer adjacent motif (PAM) and exhibits exceptionally high on-target specificity relative to other CAST systems (3).

To confirm the integration behavior of the system used in this study, we performed genome-wide profiling of transposition products recovered from *in vivo* integration assays. Integration events were highly enriched at the programmed target site, with approximately 98% of insertions occurring on target (**Figure 1B**). Analysis of insertion positions revealed a narrow integration window centered approximately 82 bp downstream of the PAM, consistent with previous observations for type I-F3 CASTs (**Figure 1C**) (3). These results establish the high fidelity and positional precision of *Vch*CAST and provide the functional context for subsequent structural investigations of the integration machinery. To facilitate cryo-EM structure determination, we employed protein engineering strategies, which will be described in future versions of this work. Throughout the manuscript, engineered proteins are referred to by their corresponding wild-type names for simplicity. The structural features, intermolecular interactions, and mechanistic conclusions reported here are independent of the specific engineering strategy employed.

### Conformational changes in the strand-transfer complex involve TnsB active site reorganization

We next determined the cryo-EM structure of the strand-transfer complex (STC), containing Integration Host Factor (IHF) (*17*), TnsA, TnsB, and a strand-transfer DNA substrate (2.9 Å overall resolution, **Figure 1D-I**, **Supplemental Figure 5**-**6**), which we refer to as Class 1. We also identified a second, minor 3D class (27% of particles, 3.2 Å overall resolution), that lacks observable density for target DNA and for TnsB beyond the C Terminal Helix (CTH) domain (Class 2, **Supplemental Figure 5, Table 1**). Closer inspection of Class 2 reveals that the catalytic DDE triad (D223, D308, and E344) is too distant (∼8 Å) to form a functional active site, and that the loop preceding E344 (residues 331–341) is disordered (**Figure 1G**). In contrast, Class 1 has the expected arrangement of catalytic residues, as well as two magnesium ions: a catalytic magnesium with the expected coordination geometry and a second structural magnesium positioned nearby that stabilizes a sharp bend at the target–donor DNA junction (**Figure 1H**). The conformational transition from Class 2 to Class 1 would entail a ∼20° rotation of TnsB’s catalytic domain (CD) and target DNA-binding domain (TBD) at the TnsA–TnsB interface (blue star, **Figure 1F**), reorienting these domains to engage target DNA. Concurrently, a target DNA-binding loop in TnsB’s catalytic domain (residues 359–378) becomes structured, further priming the complex for target DNA binding (**Figure 1I**). These structural observations show that the STC exhibits conformational heterogeneity; one conformation (Class 2) appearing catalytically inactive, and the other (Class 1) showing rearrangements consistent with active site organization and target DNA engagement. Since TnsA/B requires additional transposon-encoded partner proteins (i.e. TnsC) for integration activity (*3*), these conformational changes may reflect a mechanism of activation for TnsA/B.

### Architecture of the VchCAST holo integration complex

We next determined the cryo-EM structure of the holo integration complex (HIC), comprising Cascade-TniQ, TnsA, TnsB, and TnsC, assembled on a strand-transfer DNA substrate mimicking the *in vivo* integration product (**Supplemental Figure 6**). Our cryo-EM data analysis reveals substantial compositional heterogeneity, with three identifiable assemblies (**Supplemental Figure 7**): Cascade-TniQ (TRC, 3 Å overall resolution), STC/TnsC (4.3 Å overall resolution), and the holo integration complex (HIC, 4.5 Å overall resolution, **Figure 2A)**. In the HIC, local resolution is highest at the Cascade core (3.7 Å) and decreases toward the STC periphery (∼7 Å; **Supplemental Figure 8, Table 1**). Notably, the overall map was of sufficient quality to interpret different structural features without the need of a composite map stitched from subtracted focused-refined regions.

**Figure 2.**
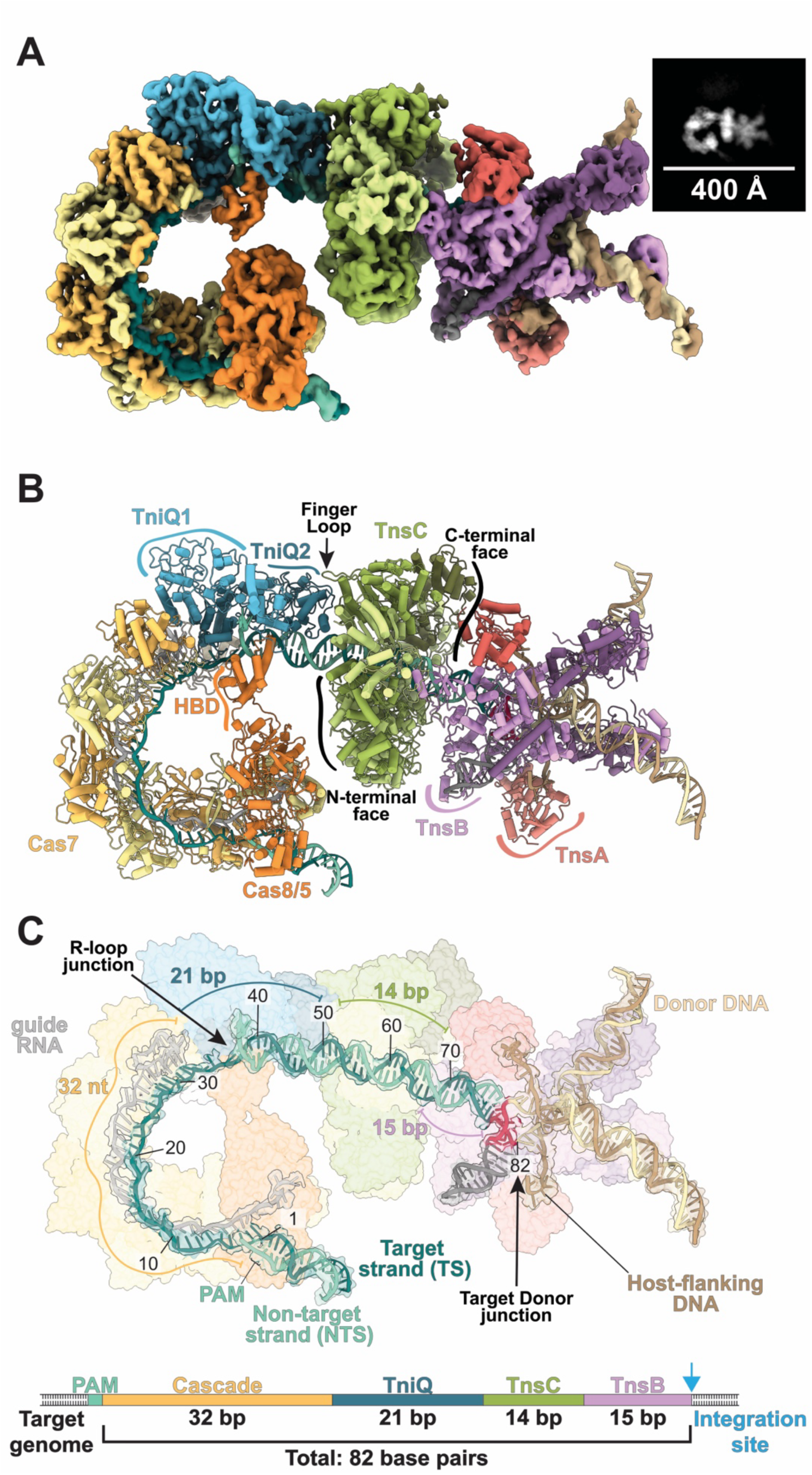
Overall architecture of the I-F3 VchCAST holo integration complex. **(A)** Cryo-EM density map of the 4.5 Å holo integration complex (HIC) displayed at a threshold of 0.077, showing all components required for integration complex assembly. Inset shows a representative 2D class average. Components colored as follows: Cas8/5, orange; Cas7, yellow; TniQ, blue, the TnsC ring, green; TnsB, purple; TnsA, red. Donor DNA is in beige and brown, guide RNA is grey, and target DNA is in teal. Alternating shades distinguish individual protomers. **(B)** Atomic model in cartoon representation. Components are labeled by protein name and structural features (HBD = helical bundle domain). The arrow points to the TnsC finger loop, and the black outline highlights the N and C terminal face of TnsC. **(C).** Surface representation of the protein components with transparency enabled. Nucleic acids are shown in cartoon representation, with segmented cryo-EM density (transparent, gray) displayed around DNA. The nucleic acid model depicts all key sequence features: PAM, target donor junction (i.e. integration site), target-site duplication (red), genomic DNA (gray), and host flanking DNA. Nucleotide positions are numbered relative to the PAM. Colored lines indicate protein footprints, and colored numbers indicate the corresponding nucleotide footprints. Cascade, TniQ, TnsC and TnsB collectively occupy 82 base pairs, consistent with the observed insertion site. Bottom: Genomic DNA-binding footprint. The blue arrow indicates the integration site. nt = nucleotide; bp = base pair.

The HIC contains a Cascade effector, a TniQ dimer, a heptameric TnsC ring, a TnsB tetramer, and two copies of TnsA (**Figure 2B**), consistent with the stoichiometry established by structures of individual subcomplexes (*10*, *11*, *13*). However, density for the seventh TnsC protomer (TnsC7) is noticeably weaker than for the other subunits (**Supplemental Figure 9**), indicating substoichiometric occupancy or flexibility. The DNA-binding footprint accounts for the characteristic 82 bp integration distance from the PAM: 32 nucleotides occupied by the guide RNA spacer, followed by a 21, 14, and 15 bp footprint corresponding to TniQ, TnsC, and TnsB, respectively, and terminating in a 5 nucleotide target-site duplication (**Figure 2C**). Mutagenesis of Cas8/5 PAM-interacting residues S127, R243, N246, and D461 substantially reduces integration activity (**Supplemental Figure 10**), confirming the importance of PAM recognition for transposition.

Our structure of the VchCAST HIC reveals an arrangement wherein TnsC acts as a bifunctional adaptor, interacting with TniQ via its N-terminal face and with the STC via its C-terminal face (**Figure 2B)**. This architectural role of TnsC is similar to what was previously observed for ShCAST (type V-K) (*18*) and PmcCAST (type I-B2) systems (*19*, *20*). This modularity provides a compelling explanation for how CAST systems have diversified to co-opt a wide range of CRISPR effectors while preserving the core transposition mechanism.

### A conserved TniQ–TnsC interface is critical for VchCAST transposition

A central mechanistic question is how the targeting module is physically coupled to the core transposition machinery. In the HIC, TniQ2 serves as the primary physical bridge between the target recognition and transposition machinery (**Figure 3A**), forming predominantly hydrophobic interactions with TnsC1 (**Figure 3B**). Consistent with the functional importance of this interface, alanine substitutions at F2, L3, and F40 significantly reduced integration activity (**Figure 3C**). Although TniQ forms a constitutive homodimer, only the TniQ2 monomer interacts with TnsC. This parallels the evolutionarily distant ShCAST and PmcCAST, in which a single TniQ subunit recruits TnsC (*18*, *19*). Remarkably, despite the divergent overall architectures of the ShCAST (PDB: 8EA3), PmcCAST (PDB: 8FF4), and VchCAST integration complexes, the TniQ–TnsC interface is strikingly conserved (**Supplemental Figure 11**). In particular, the TnsC finger loop, previously shown to mediate critical TniQ interactions in ShCAST (*18*), engages TniQ2 similarly in VchCAST (**Supplemental Figure 11**). Consistent with this observation, truncations within the TnsC finger loop severely reduces transposition activity (*10*). Together, these findings identify the TniQ-TnsC interface as a shared docking point that may enable architecturally diverse CRISPR effectors to couple to common transposition machinery.

**Figure 3.**
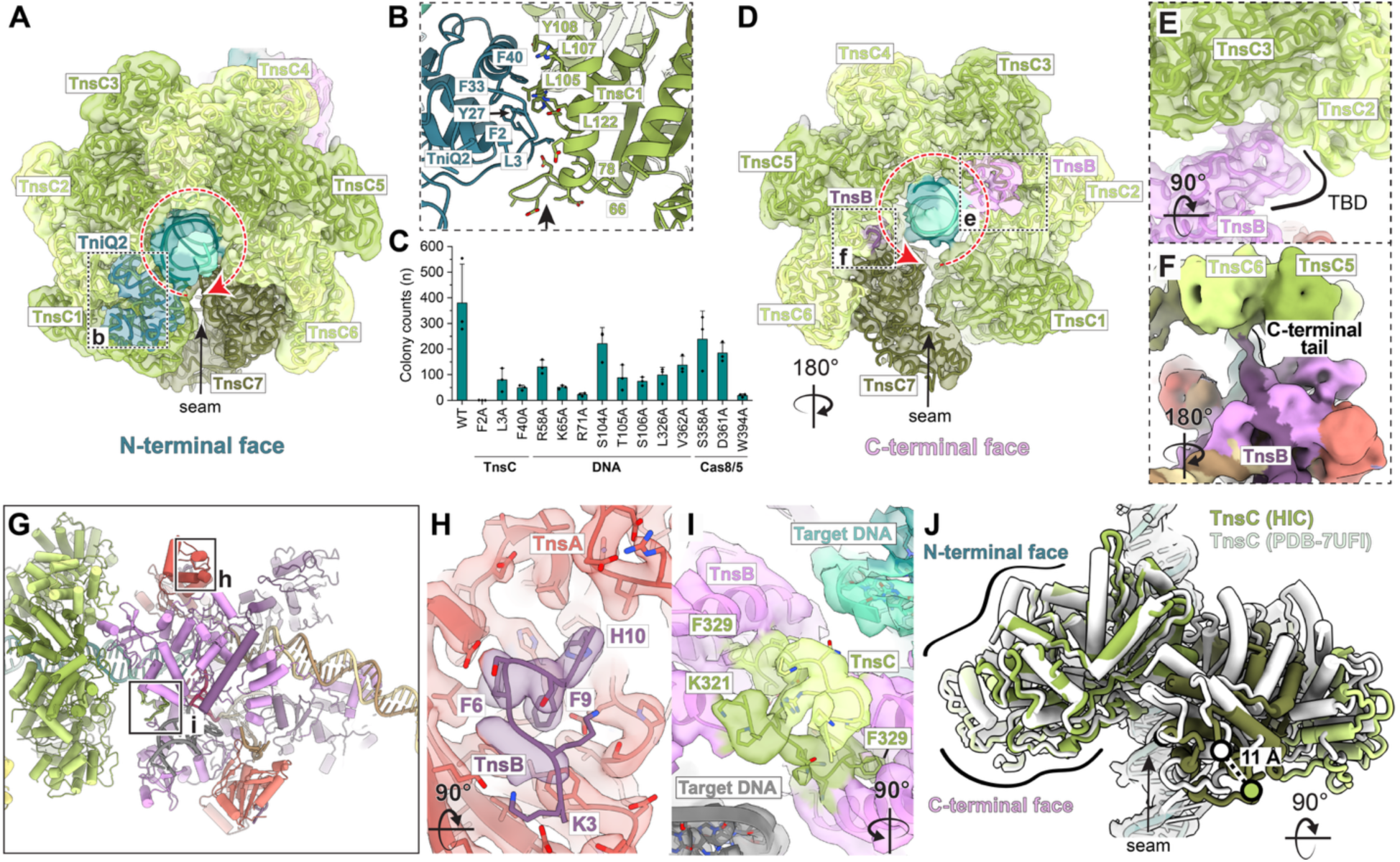
Critical protein-protein interactions within the holo integration complex. **(A)** N-terminal face of the TnsC heptamer. The atomic model is shown within the transparent HIC cryo-EM density map (threshold = 0.077). TnsC subunits are numbered beginning with the protomer that interacts with TniQ2. The red arrow indicates the sequential arrangement of TnsC protomers forming heptameric ring, and the black arrow points to the TnsC seam. The boxed region highlights the TniQ2-TnsC1 interface shown in panel B. **(B)** Molecular details of the TniQ2-TnsC1 interface. The arrow points to the TnsC finger loop. Positions 66 and 78 are indicated to show the beginning and end of the TnsC finger loop. Key residues mediating TniQ2-TnsC1 interactions are shown as sticks and labeled using the one-letter amino acid code and residue position. **(C)** Effects of TniQ mutations measured by *in vivo* transposition assays. The y-axis represents colony counts after kanamycin selection (overall activity). WT = wild-type TniQ. The assays were performed in biological triplicate. Mean ± standard deviation shown; n=3 per bar. Residues are grouped by category: TnsC, residues at the TniQ2-TnsC1 interface; DNA, DNA interacting residues; Cas8/5, residues at the TniQ-Cas8/5 interface. **(D)** C-terminal face of the TnsC heptamer. The atomic model is shown within the transparent HIC cryo-EM map using display parameters identical to those in panel A. The red arrow indicates the TnsC protomer order following the convention established in panel A. Boxed regions indicate TnsB-TnsC interactions shown in panels E and F. The view is rotated relative to panel A. **(E)** Detailed view of TnsB interactions spanning the TnsC inter-protomer interface. The HIC cryo-EM density map is shown as a transparent surface using display parameters identical to those in panels A and D. The atomic model is shown in cartoon representation. Coloring and labeling follow the conventions established in panels A and D. TBD = TnsB target-binding domain. The rotation is relative to panel D. **(F)** Low-pass filtered (10 Å) HIC cryo-EM density revealing the flexible TnsB C-terminal tail. Rotation is relative to panel E. **(G)** HIC model with docked C- and N-termini of TnsC and TnsB from the STC/TnsC and STC structures, respectively. Black boxes highlight the interactions shown in panels H (TnsA–TnsB) and I (TnsB–TnsC). Disordered tails are shown as thick cartoons for visual clarity. **(H)** Interactions between the TnsB N-terminus (residues 3-10) and TnsA. The atomic model (cartoon representation) is docked into transparent STC cryo-EM density (Class 1, threshold = 0.11). Residues are shown as sticks, and selected residues are labeled using one-letter amino acid code and residue number. The view is rotated 90° relative to panel G. **(I)** Interactions between the TnsC C-terminal tail and TnsB. The atomic model is shown within transparent 4.3 Å STC/TnsC cryo-EM density (threshold = 0.06). Selected TnsC residues are labeled. The view is rotated 90° relative to panel G. **(J)** Comparison of the TnsC oligomeric structure in the HIC and a previously determined DNA-bound TnsC heptamer (PDB-7UFI, white), aligned on TnsC1. The view is rotated 90° relative to panel A. The 11 Å displacement of TnsC7 is indicated by circles (white = 7UFI, green = HIC) and the measured distance is shown with a dotted line.

### The holo integration complex reveals critical interactions between TnsB and TnsC

TnsB activity is strictly dependent on TnsC-mediated recruitment to the target site (*10*), making these interactions central to understanding integration. We next examined how TnsB engages TnsC within the HIC, revealing multiple points of contact along TnsC’s C-terminal face, specifically at protomers TnsC2, TnsC3, and TnsC6 (**Figure 3D**). Notably, TnsB’s TBD localizes to the TnsC2–TnsC3 interface (**Figure 3E**), consistent with previous ShCAST and PmcCAST structures (*18*, *20*) (**Supplemental Figure 12**). Across Tn7-like transposons, ATP binding, but not hydrolysis, is required for integration activity (*21*, *22*). AAA+ initiator clade regulators require ATP for oligomerization, as the ATP-binding pocket is formed at the inter-protomer interface (*23*). This implies that transposition requires TnsC oligomerization on target DNA, though the mechanistic basis for this requirement has remained unclear. The conserved interaction between TnsB’s TBD and the TnsC inter-protomer interface now suggests that TnsB directly senses the oligomeric state of TnsC, providing a structural explanation for this long-standing observation.

Another conserved feature across CAST systems is the interaction between TnsC and the extreme C-terminus of TnsB, known as the hook, which is critical for transposition (*24*). The interchangeability of TnsB hooks between type I-F3 CASTs (*25*) further underscores the importance of these structured interactions within otherwise disordered regions. Consistent with this, we observe continuous density connecting the CTH (residues 563-573) of TnsB to the surface of TnsC6 (**Figure 3F**), though additional contacts likely go undetected due to the length and extensive disorder of this region. The TnsB hook is predicted by AlphaFold3 (*14*) to correspond to the last 8 residues of TnsB, an assignment supported by cryo-EM density observed in the HIC (**Supplemental Figure 13**). The hook binding site in VchTnsC aligns well with ShTnsC (*24*) and PmcTnsC (*20*) (**Supplemental Figure 13**), indicating that although structural details may differ, the binding location is conserved.

### Core transposition machinery makes extensive intermolecular interactions via disordered termini

Having confirmed the TnsB hook interaction with TnsC, we reasoned that the disordered N- and C-terminal regions present in all components of the core transposition machinery (TnsABC; **Supplemental Figure 14**) likely mediate additional structured interactions. Consistent with this, we observe contacts between the N-terminus of TnsB with TnsA, and between the C-terminal tail of TnsC with TnsB (**Figure 3G**). Residues 3–10 of TnsB’s disordered N-terminus occupy a hydrophobic pocket on TnsA (**Figure 3H**) through aromatic residues F6 and F9, consistent with AlphaFold3 predictions (*14*) (**Supplemental Figure 1**) and sequence conservation analysis (**Supplemental Figure 15**). Similarly, the TnsC C-terminal tail (residues 321–330) forms hydrophobic interactions with the TnsB C-terminal helix (CTH; **Figure 3I**), also supported by AlphaFold3 predictions (*14*) (**Supplemental Figure 16**) and conservation analysis (**Supplemental Figure 15**). Truncations of TnsC residues 304–330 result in complete loss of transposition activity (*10*), supporting the importance of this structured interaction. Analogous interactions between the TnsC C-terminus and the STC have been observed in PmcCAST (*20*) (**Supplemental Figure 17**). Together, these findings uncover a previously unrecognized network of interactions at the unstructured termini of TnsA, TnsB, and TnsC, revealing how the core transposition machinery may maintain associations to facilitate integration competent assembly.

### The structure of the TnsC heptamer is influenced by context

A previous cryo-EM structure of VchTnsC solved in the presence of DNA revealed an open heptameric ring-like arrangement in which the bound DNA was poorly resolved (PDB-7UFI) (*10*). In our HIC structure, although we clearly resolved the path of target DNA, no specific TnsC-DNA interactions could be precisely mapped. Thus, it is likely that the TnsC ring primarily associates with target DNA via the complementary basic surface formed by its central pore (**Supplemental Figure 18**). This is consistent with previous data showing that TnsC binds DNA relatively weakly (*10*), suggesting that TnsC requires partner proteins like TniQ to initiate oligomerization on target DNA. Strikingly, comparison with PDB-7UFI reveals that the heptameric ring is significantly more open in the HIC. Aligning on TnsC1 results in an 11 Å displacement at TnsC7 relative to the equivalent protomer in PDB-7UFI (**Figure 3J**), the result of an average 0.8 Å greater rise per protomer (**Supplemental Figure 19**).This indicates that partner proteins induce more dramatic conformational changes than those observed in TnsC with DNA alone. Altogether, our observations suggest that TnsC’s oligomeric state is sensitive to the presence or absence of specific binding partners and is mediated through distinct interaction interfaces on specific TnsC protomers.

### Concerted structural remodeling of Cascade and TniQ license TnsC recruitment

We were particularly interested to know how TnsC is selectively recruited to the target site. The compositional heterogeneity of our reconstitution allowed us to resolve a free Cascade–TniQ complex (**Supplemental Figure 7**) named here as the target–recognition complex (TRC, 3.0 Å overall resolution; **Figure 4A**), closely resembling a prior structure of the target DNA bound VchCAST Cascade-TniQ complex (*11*) (**Supplemental Figure 20**). This previously reported state featured a partial R-loop and a disordered Cas8/5 helical bundle domain (HBD) (*11*).In contrast, the Cascade–TniQ complex in the HIC (**Figure 4B**), adopts a fully engaged conformation characterized by a complete R-loop and a structured Cas8/5 HBD. Comparison of the Cascade–TniQ in the TRC and HIC structures reveals that R-loop completion is accompanied by a conformational change in TniQ2 to engage both target DNA and TnsC (**Supplemental Figure 21**).

**Figure 4.**
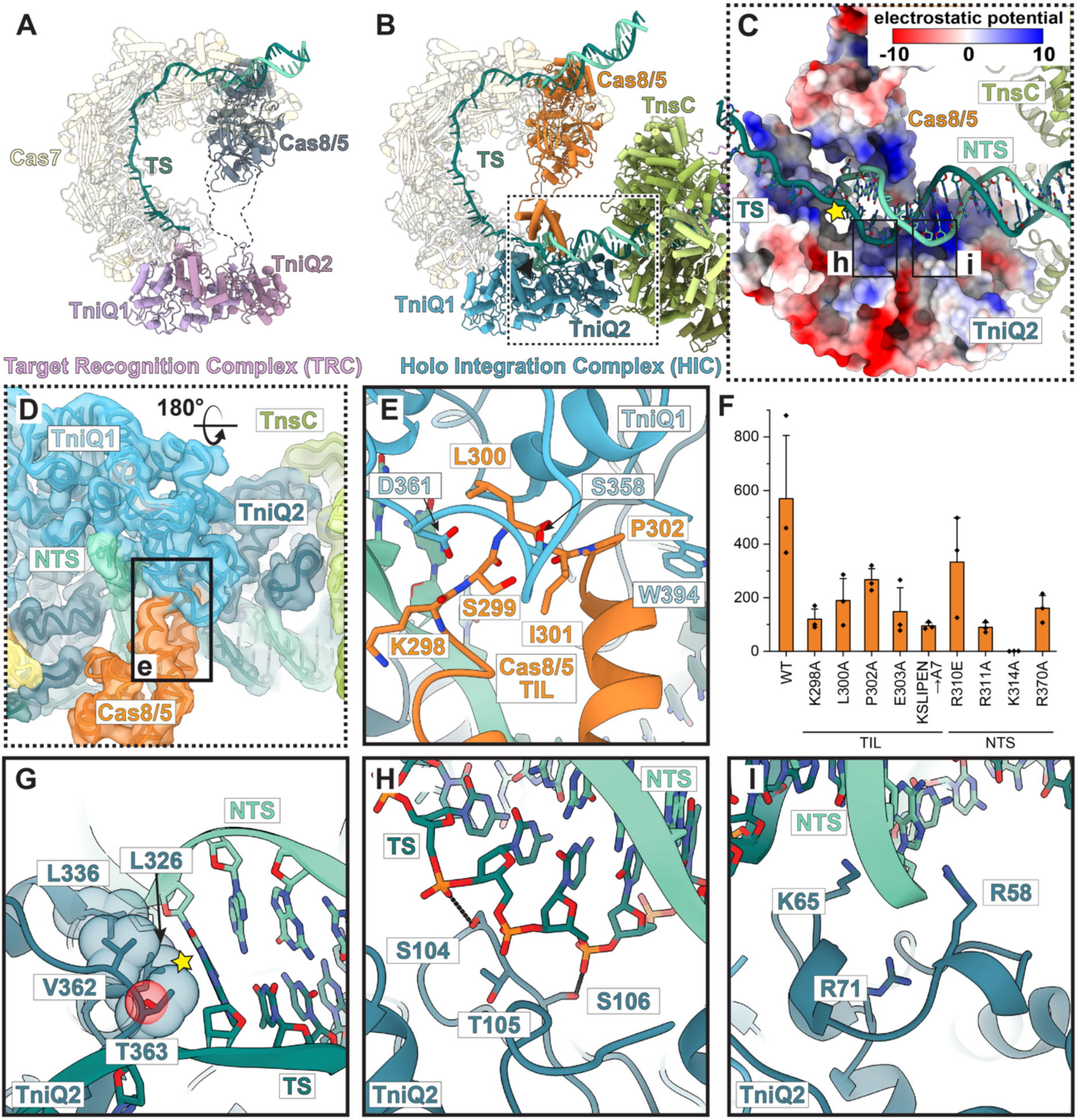
Cascade remodeling establishes a Cas8/5–TniQ junction that recognizes the R-loop and licenses recruitment of TnsC. **(A)** Atomic model of the target recognition complex (TRC), showing the unstructured Cas8/5 helical bundle domain (HBD, indicated by dotted lines), resulting in a partially unstructured target DNA. TS = target strand (teal). Individual copies of TniQ are numbered as TniQ1 (light pink) and TniQ2 (dark pink); Cas8/5 (gray) and Cas7 (transparent) are labeled. **(B)** Atomic model of the holo integration complex (HIC), focusing on the Cascade-TniQ region. This transition (TRC to HIC) orders the target DNA duplex and facilitates recruitment of the TnsC ring. The boxed region (dotted lines) indicates the areas enlarged in C, D, and E. **(C)** A positively charged surface at the Cas8/5–TniQ1 junction forms the binding interface for both the target strand (TS) and non-target strand (NTS). Surfaces are colored according to electrostatic potential (legend, upper right). Positive values reflect positive coulombic charge, and negative values reflect negative coulombic charge. The yellow star indicates the R-loop junction and the view shown in panel G. Lowercase lettered boxes indicate the views shown in panels H and I. **(D)** The narrow channel formed by TniQ1, TniQ2, and Cas8/5 accommodates threading of the non-target DNA strand. The atomic model is shown fitted into the cryo-EM density map (transparent HIC surface, threshold = 0.077). The boxed region corresponds to the view shown in panel E. Rotation indicated at the top of the panel is relative to panel B. **(E)** The Cas8/5 helical bundle domain loop (named as TniQ-interacting loop (TIL), residues 298–302) interacts with TniQ1. Residues mediating Cas8/5 - TniQ interactions are shown as sticks and labeled using one-letter amino acid code and residue number. The view is identical to that shown in panel D. **(F)** Effects of Cas8/5 mutations measured by *in vivo* transposition assays. The y-axis represents colony counts after Kan selection (overall activity). WT = wild-type Cas8/5. The assays were performed in biological triplicate. Mean ± standard deviation is shown; n=3 for each bar. Residues are grouped by category: TIL, TniQ-interacting residues in Cas8/5, NTS, Cas8/5 residues interacting with the non-target strand. **(G)** Selected TniQ2 residues interacting with the target DNA duplex at the R-loop junction (yellow star). Residues are labeled and shown as transparent spheres and sticks. **(H-I)** Molecular details of the view diagramed in panel C, showing interactions between TniQ2 and target DNA. Putative hydrogen bonding interactions (black dashed lines) are shown between TniQ2 residues S104 and S106, shown as sticks, and the target DNA backbone.

Strikingly, the non-target strand (NTS) is threaded through an electropositive channel at the tripartite TniQ1–TniQ2–Cas8/5 HBD interface, where it rejoins the target strand (TS) to reform the target DNA duplex downstream of the R-loop junction (**Figure 4C & D**). The Cas8/5 HBD contributes to this channel through electrostatic contacts with the NTS via R310, R311, K314, and R370 (**Supplemental Figure 22**). Additionally, a previously uncharacterized loop within the Cas8/5 HBD (residues KSLIPEN), which we term the TniQ–interacting loop (TIL), directly interacts with TniQ1 (**Figure 4E**). The functional importance of this interface is confirmed by alanine substitutions within the Cas8/5 HBD, the TIL, and at TniQ1 residues D361, S358, and W394, all of which substantially reduce integration activity (**Figures 3C & 4F**).

Separately, TniQ2 engages target DNA through two distinct modes: hydrophobic packing against the R-loop junction (L326, L336, V362, and T363; **Figure 4G**) and electrostatic interactions with the DNA backbone via polar and basic residues (R58, K65, R71, S104, and S106; **Figure 4H-I**). Mutagenesis confirms the importance of both sets of interactions (**Figure 3C**). Collectively, these observations establish that R-loop completion drives a conformational transition in the target recognition module that repositions TniQ to engage target DNA and license recruitment of the core transposition machinery.

## Discussion

Our findings lead us to propose the following working model for RNA-guided transposition, incorporating prior observations on CASTs and other Tn7-like systems. Cascade-TniQ promiscuously samples genomic locations (*10*), consistent with its ability to stably bind target DNA with only partial R-loop formation (**Figure 5**). A cooperative conformational rearrangement in Cas8/5 and TniQ is required to stabilize the full R-loop and enable TnsC recruitment, with TniQ’s direct contacts at the R-loop junction providing a structural readout of R-loop completion. This coupling is further supported by molecular dynamics simulations demonstrating correlated motions between Cas8/5 and TniQ (*26*). Beyond this, we observe no DNA distortions consistent with specific recognition of the target site by the transposon-encoded machinery (**Supplemental Figure 23**), contrary to that observed in other Tn7-like systems (*20*, *27*). Further insight into target-site selectivity comes from the observation that PAM-distal mismatches permit Cascade binding but abolish transposition (*10*). The conformational rearrangements in the HIC occur near spacer nucleotide positions 25–32, raising the possibility that mismatches in this region could introduce geometric incompatibility that prevents stable docking of the helical bundle domain (HBD) and subsequent TnsC recruitment (**Figure 5**). TnsC oligomerization is required to recruit TnsA/B to the target site and may be sensed directly by TnsB through conserved interactions spanning TnsC protomers. Whether full heptamer formation is required prior to TnsA/B recruitment remains unknown and is not resolvable using our reconstitution approach; however, previous work on Tn7 suggests that oligomerization is completed prior to TnsA/B recruitment (*28*).

**Figure 5.**
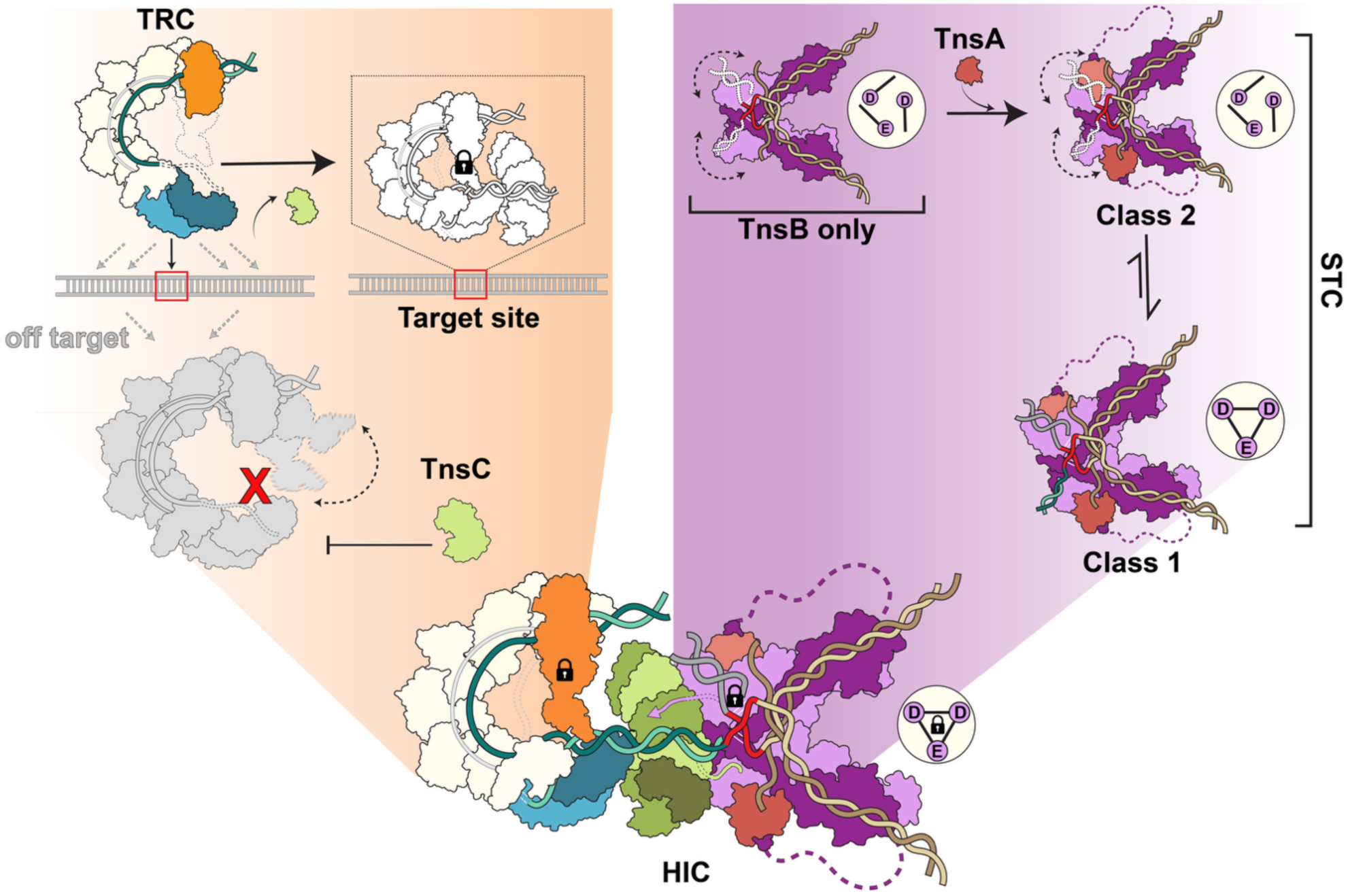
Proposed licensing mechanisms regulating type I-F3 VchCAST assembly. Two major licensing events occur during RNA-guided DNA integration: 1. at the CRISPR effector (orange box) and 2. at the transposase (purple box). **(A)** Orange box: Top left, the target recognition complex (TRC) initially displays an unstructured Cas8/5 helical bundle domain, a partial R-loop, and outward-open TniQ conformation. Unstructured regions are indicated with dotted lines. This state can sample many genomic locations (dotted arrows). The red box indicates the programmed target site. Top right, upon recognition of the correct target, the TRC forms stabilizing interactions between Cas8/5 and TniQ that promote R-loop completion and enable recruitment of TnsC. The completion of this licensing step is represented by the lock symbol. Middle, off-target complexes (gray silhouette) are unable to form a complete R-loop (red X) and the Cas8/5 helical bundle domain remains flexible (bi-directional black arrow). As a result, TnsC (green) cannot be recruited. **(B)** Purple box, top left: The TnsB-only strand transfer complex (STC) contains flexible target-binding domains (bi-directional arrows), and an incompletely arranged active site, with catalytic triad DDE residues shown as disconnected lines inside white circle. Top right and middle: upon addition of TnsA, the TnsA/B STC adopts two conformational states that exist in equilibrium: Class 2 and Class 1. Class 1 exhibits the catalytically competent arrangement of the DDE triad (indicated by the triangle within the circle). Dotted lines indicate flexible connections between components, including the interface between TnsA and TnsB. **(C)** The targeting module (orange) and transposition module (purple) assemble with TnsC to form the holo integration complex (HIC). The two lock symbols indicate that both the targeting and the transposition module have adopted stable conformations and are locked into an integration competent conformation. A third lock symbol on the DDE triad denotes formation of the catalytically competent active site. The TnsB and TnsC C-terminal tails function to loosely couple components, with the TnsB ‘hook’ inserting into a TnsC pocket, forming β-sheet interactions (purple arrow), while the TnsC C-terminal tail contacts TnsB near target DNA. Tail colors match their parent proteins, and dotted lines indicating disordered regions.

TnsB also undergoes multiple layers of activation through association with partner proteins TnsA and TnsC. In isolation, the DDE catalytic triad is improperly arranged and too distant to coordinate magnesium ion binding, and target DNA remains disordered even when a strand-transfer substrate is provided (*13*). Association with TnsA is insufficient to fully stabilize the catalytically competent conformation (**Figure 5**). Only in the context of the holo integration complex, where both TnsA and TnsC are present, the catalytic residues are locked into an integration-competent state and target DNA is stably accommodated. Similar activation mechanisms have been previously observed for the IS21 elements, in which IstB creates a conformational change in IstA, the transposase, activating it for DNA-integration (*29*). Thus, conformational changes appear to be a highly effective mechanism for ensuring selective activation of integration. In summary, we propose that each component of the integration machinery undergoes conformational changes upon incorporation into the holo integration complex, providing a structural basis for the exceptionally high on-target specificity characteristic of type I-F3 CAST systems. Both the targeting module, Cascade–TniQ, and the transposase, TnsA and TnsB, serve to license RNA-guided DNA integration, ensuring that catalytic competence is only achieved upon assembly of the complete machinery at the RNA-guided target site.

Structured interactions embedded within intrinsically disordered regions of CAST proteins have been shown to play critical roles in transposition, most notably through the TnsB C-terminal hook (*18*, *30*). Here, combining our cryo-EM structure with AlphaFold3 predictions, we precisely map the TnsB C-terminal hook and confirm its binding pocket on TnsC. Beyond this, we identify two previously unrecognized interactions of this type: the TnsB N-terminus engaging a pocket on TnsA, and the TnsC C-terminus contacting TnsB in close proximity to target DNA. Together, these findings reveal an unexpectedly extensive network of structured interactions within otherwise disordered termini that physically couple the core transposition machinery, likely forming a loose network of interactions between TnsA, TnsB, and TnsC prior to integration complex assembly. The precise mapping of these contacts provided here has direct implications for the rational engineering of chimeric CAST systems and the design of improved variants.

In closing, this work establishes a comprehensive mechanistic framework for RNA-guided DNA integration that appears to hold across CAST diversity. With a high-resolution structure of the holo integration complex now in hand, we anticipate that continued structure-guided optimization will drive CAST systems toward the efficiency and precision required for broad translational applications.

## Acknowledgements

We acknowledge Ines Chen, Alexandre Zanghellini, Tuba Şevik, and the rest of the Kellogg lab for helpful discussions related to cryo-EM imaging, analysis, and manuscript feedback. We also acknowledge Asfarul Haque for exceptional support of cryo-EM imaging during this time. This research is supported by the Pew Charitable Trusts (E.H.K.), NIH NIGMS 7R01GM144566 (E.H.K.), NIH NIGMS F31GM151863 (V.H.T.), and the Cystic Fibrosis Foundation (E.H.K.). We also thank the St. Jude Hartwell Center for Bioinformatics & Biotechnology which is supported in part by ALSAC and the National Cancer Institute grant P30 CA021765 for their help conducting next-generation sequencing, the St. Jude Cryo-EM Center, and St. Jude Research Information Services for providing high-performance scientific computing systems and computational resources that contributed to the research in this manuscript. The authors used Claude (Anthropic) to assist with manuscript editing and language refinement. All scientific content, ideas, and interpretations are solely the work of the authors. The funders had no role in study design, data collection and analysis, decision to publish, or preparation of the manuscript.

